# Latent-Factor Stability Is Not Sufficient: Evaluating SABLE-MO in Simulated and Cancer Multi-Omics Data

**DOI:** 10.64898/2026.10.12.758631

**Authors:** Elena Krikun, Abedalrhman Alkhateeb

## Abstract

Reproducibility is often used to assess latent factors learned from multi-omics data. Yet factors that recur across model refits may not reflect the shared biological structure of interest. We investigated this issue using SABLE-MO, a shared/private representation-learning framework with a separate factor-retention procedure. We evaluated the method in simulations with known latent structure. The original retention rule produced no false biological shared-factor acceptance across 800 independent-null datasets, but accepted at least one false factor in 70 of 200 correlated-private datasets (35%). Following an exploratory analysis of this failure, we increased the lower-decile aligned-correlation threshold from 0.40 to 0.60 and tested it on a new, independently seeded simulation bank. False acceptance decreased from 33% to 19%, while recovery of true shared factors changed only slightly (94.375% to 94.0%). Exact modality-support attribution reached 67.5%, below its prespecified criterion. We also analyzed matched RNA expression, DNA methylation, and copy-number data from TARGET osteosarcoma (78 patients), TCGA-SARC (250), and TCGA-KIRC (304). Outcome-blind model selection favored a simpler shared/private representation in TARGET. In both TCGA cohorts, the more complex representation improved cross-view retrieval and missing-view reconstruction relative to the simpler model, but worsened ordinary reconstruction. These external comparisons were performed before factor retention. Our results show that factor reproducibility, representation quality, and biological interpretation require separate assessment. Stricter retention reduced errors in a specific simulated setting but did not eliminate them.

## I. Introduction

Multi-omics analysis integrates measurements such as RNA expression, DNA methylation, and copy number from the same samples. A shared representation may summarize joint patterns, but the presence of a reproducible latent direction does not establish that it corresponds to a biological factor of interest. Correlated modality-specific variation or technical structure can also recur across refits. Conversely, two equivalent fits may recover the same subspace with different rotated axes.

The distinction between joint and individual structure is already established in JIVE and AJIVE [1], [2]. MOFA offers a probabilistic alternative for latent multi-omics integration [3]. Studies of multi-omics reproducibility and null behavior [4], as well as stability selection [5], motivate systematic evaluation, but do not furnish an error-control guarantee for the particular factor-retention rule studied here. Rather than claiming a fundamentally new shared/private architecture, we examine *when its stable factors are admitted for biological interpretation*.

We investigate three linked questions. First, does a prespecified retention rule falsely accept shared biological factors in a structured null retaining cross-view dependence? Second, can one frozen modification reduce that error on new simulation data without a large recovery loss? Third, what representation-level evidence exists in three cancer cohorts, where the simulated latent truth is unavailable? The principal contribution is a mechanism-specific operating-characteristic analysis with a separately tested correction. The real-cohort comparisons are deliberately not described as validation of the complete retention gate.

## II. Methods

### A. Shared/private representation learning

For *V* = 3 matched assay blocks 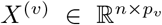, the SABLE-MO representation engine uses modality-specific encoders, a shared patient representation formed by averaging available-view shared scores, and view-private scores. The compact selected configuration uses five shared dimensions, two private dimensions per view, and 32-unit hidden layers. Its objective combines block-normalized reconstruction, cross-view alignment, contrastive matching, orthogonality/separation penalties, and missing-view reconstruction; modality dropout simulates absent assays during training. These component families have precedents in multi-view learning and modern neural optimization [6]–[8].

Within each training or resampling fit, filtering, median imputation, standardization, and modality-specific PCA are refitted rather than transported from the full dataset. The fixed feature budgets are at most 5,000 RNA-like, 10,000 methylation-like, and 5,000 copy-number-like features, followed by at most ten PCA dimensions per block. The frozen simulation model uses full-batch AdamW (learning rate 10^−3^, weight decay 10^−4^), with 248 epochs for each historical-gate and prospective-validation reference/subsample fit. These parameters and all retention rules were fixed before their designated confirmatory evaluation.

### B. Prespecified factor retention

For each dataset, the reference representation is evaluated against *B* = 100 complete pipeline refits on 80% subsamples. Latent spaces are aligned using orthogonal Procrustes [9], and shared factors are paired by Hungarian matching [10]. Let *r*_*kb*_ be the absolute correlation for factor *k* against matched refit *b*. A match is counted present when *r*_*kb*_ ≥ 0.50, yielding *π*_*k*_ = *B*^−1^ ∑ _*b*_ **1***{r*_*kb*_ ≥ 0.50*}*. Decoder-weight energies determine supported views: a modality supports factor *k* if its normalized first-layer decoder-weight energy for *k* is at least 0.15.

A factor is retained only if *all five* conditions in Table I hold. Thus the per-view 0.15 energy requirement defines the support count; it is not a sixth independent Boolean criterion. Axis-level retention, rotational-subspace recovery, and exact generating-modality attribution are evaluated separately. The energy definition is tied to the frozen model parameterization and is not a general explanation of feature importance.

**TABLE I.** Frozen FACTOR-RETENTION RULE. A SINGLE LOWER-TAIL THRESHOLD CHANGES IN THE FOLLOW-UP VALIDATION; THE OTHER CRITERIA REMAIN FIXED.

| Criterion | Original | Candidate |
| --- | --- | --- |
| Median aligned correlation | $\geq 0.70$ | $\geq 0.70$ |
| Lower-decile correlation, $q_{0.10}$ | $\geq 0.40$ | $\geq 0.60$ |
| Inclusion frequency, $\pi_k$ | $\geq 0.80$ | $\geq 0.80$ |
| Factor score variance | $\geq 10^{-4}$ | $\geq 10^{-4}$ |
| Supported modalities | $\geq 2$ | $\geq 2$ |
| Defining support: relative energy | $\geq 0.15$ | $\geq 0.15$ |

### C. Known-truth simulation and structured nulls

Simulation datasets distinguish generating biological-shared scores *S*, private scores *P* ^(*v*)^, technical scores *T*, and noise *E*^(*v*)^:

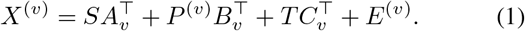

For the correlated-private null (*n* = 62), there are no *generator-designated* shared biological factors. However, private scores share a Gaussian component,

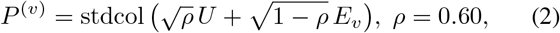

with per-column standardization before and after mixing. Hence the null does *not* mean independent modalities, and *ρ* is a nominal generator parameter rather than each dataset’s realized cross-view correlation. A false acceptance is a *dataset-level* event: at least one retained factor matches no registered biological-shared truth, subject to the prespecified matching and technical-flag protocol. Rates are conditional on the simulation truth taxonomy, not universal error guarantees. For standard-signal datasets, recovery instead counts truth-factor opportunities; support-attribution accuracy has a third, conditional denominator.

The historical 24-scenario, 3,100-dataset registry contains independent, correlated-private, standard-signal, rotational, mixed-support, and technical-null conditions. Its frozen core evaluation is not retroactively changed by the follow-up. Post-result exploration examined ablations, factor-level diagnostics, and candidate thresholds. One threshold (*q*_0.10_ = 0.60) was then locked before a new 400-dataset bank with non-overlapping random seeds (200 correlated-private; 200 standard-signal datasets, four true factors each). Original and candidate retention were applied to *the same fitted summaries* in this fresh bank, permitting paired comparison. Confidence intervals use Wilson intervals for proportions [11]; the absolute paired reduction uses a dataset bootstrap, and discordance uses an exact McNemar test [12]. Simulations follow distinct development, exploration, and validation roles [13].

### D. Outcome-blind cancer-cohort applications

The real-data analysis uses 78 matched TARGET-OS patients [14], 250 TCGA-SARC patients [15], and 304 TCGA-KIRC patients [16], each with matched RNA, methylation, and copy-number measurements. These are *different patients and independent within-cohort fits*, not transfer of TARGET-trained weights to TCGA. TARGET representation selection used five repeats of five-fold outer cross-validation and inner-fold model selection based on an outcome-blind utility of held-out reconstruction (0.30), adjusted cross-view retrieval (0.30), missing-view reconstruction (0.25), and modality balance (0.15), followed by an eligible one-standard-error selection rule. The historical selection favored V1, the shared/private autoencoder, rather than V4’s more elaborate representation engine. V3 and V4 share the training architecture and objective; V4’s defining post-fit retention operation is a distinct stage.

SARC/KIRC used five outer folds and two fixed neural seeds per fold. The compared methods were budgeted block PCA (*K* = 10), V1, and the *pre-retention V4 representation*; they were not a validation of the V4 factor-retention decision. Omitted-view NMSE and ordinary reconstruction NMSE are lower-is-better, while chance-adjusted cross-view mean reciprocal rank (MRR) is higher-is-better. The external molecular-target NMSE is not on the same scale as the bounded, PCA-space utility components in TARGET. MOVE and iCluster2b were assessed in a separate simulation comparator branch, not in these external comparisons.

## III. Results

### A. Historical retention: the pooled metric concealed a specific failure

The historical pooled biological-null false-acceptance rate was 70/1,000 (7.0%, Wilson upper 95% endpoint 8.75%), meeting its prespecified pooled rule. Yet all 70 events occurred in 200 correlated-private null datasets (35%); the other 800 independent-null datasets produced zero such events. At the matched sample size *n* = 62, the separate independent-null subset had 0/200 events, versus 70/200 correlated-private events (Fig. 1). These groups differ by generating mechanism and are not paired samples. Because dependence persists in the correlated-private null, the failure cannot be reduced to a problem of unstable factor fitting alone.

**Fig. 1.**
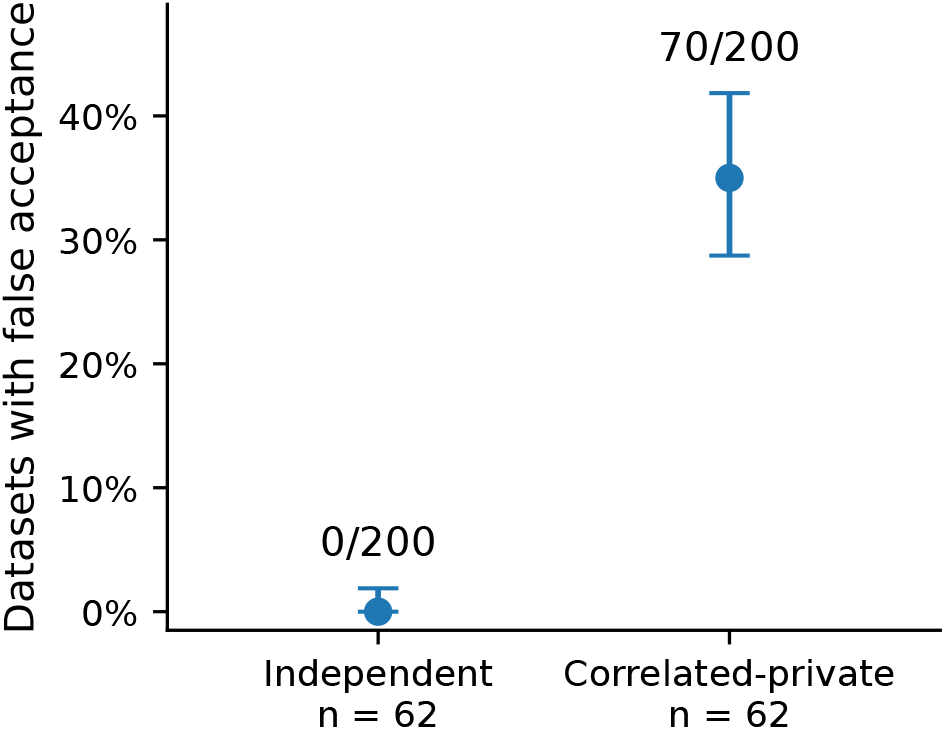
Historical matched-*n* null comparison, *n* = 62. Dataset-level false acceptance: 0/200 independent versus 70/200 correlated-private; whiskers are marginal 95% Wilson intervals. The full independent-null bank contains 800 datasets over multiple sample sizes.

Other historical endpoints used distinct populations (Table II). Standard true-factor recovery was 371/400 (92.75%). Rotational subspace recovery reached a mean canonical-correlation summary of 0.829, illustrating that subspace information can survive changing factor axes. However, exact generating-modality support was identified for only 81/120 eligible factors (67.5%), *failing* its 80% criterion. Thus a passing pooled null test did not justify unrestricted statements about the provenance or modality identity of retained biological factors.

**TABLE II.** Selected HISTORICAL FINAL-GATE OUTCOMES. They REFER TO THE ORIGINAL RULE, NOT THE LATER *q*0.10 = 0.60 CANDIDATE.

| Endpoint | Result | Interpretation |
| --- | --- | --- |
| Pooled biological-null datasets | 70/1000 | PASS |
| Independent-null datasets | 0/800 | — |
| Correlated-private null | 70/200 | Failure concentration |
| True shared-factor recovery | 371/400 | PASS |
| Rotational subspace recovery | 0.829 | PASS |
| Exact modality-support attribution | 81/120 | FAIL |
| Technical-null false acceptance | 0/400 | PASS |

### B. Diagnostics and prospective threshold validation

In archived explorations restricted to historical datasets, removing the support-count and variance requirements did not change the matched-*n* correlated-private false-acceptance count of 70/200. A grouped out-of-fold classifier using median, lower-decile, and inclusion summaries distinguished retained false factors from retained standard-signal factors with AUC 0.967 (95% bootstrap 0.950–0.981). This comparison couples scenario with truth, however, and is *not* evidence of a general biological-factor classifier. A non-retained scenario-label anchor had AUC 0.792, whereas realized private-score correlation discriminated false-acceptance events weakly (AUC 0.579, interval including 0.5). These exploratory diagnostics suggested lower-tail reproducibility was informative within this frozen simulation family, without establishing causation or identifiability.

Exploratory threshold slices showed multiple possible tradeoffs: with median 0.70 and inclusion 0.80 held fixed, *q*_0.10_ = 0.60 yielded 41/200 historical correlated-private false acceptances and 368/400 standard truth opportunities recovered; *q*_0.10_ = 0.65 yielded 18/200 and 355/400, respectively. Hence 0.60 was *not* the only admissible or optimal choice. It was selected as one documented trade-off and frozen for prospective checking.

On the new correlated-private bank, the original rule falsely accepted a factor in 66/200 datasets (33%), compared with 38/200 (19%) under the candidate (Fig. 2). The absolute *paired* reduction was 14 percentage points (dataset-bootstrap 95% CI 9.5–19.0); 28 datasets changed from false acceptance to nonacceptance and none reversed (exact two-sided McNemar *p* = 7.45 *×* 10^−9^). In a separate new standard-signal bank, true-factor recovery was 755/800 (94.375%) versus 752/800 (94.0%). The candidate satisfied the frozen directional improvement and minimum recovery conditions. The remaining 19% false acceptance is substantial; prospective validation here means fresh seeds under the same simulator, not external biological replication. The historical frozen gate and its D05 failure were unchanged. The original technical-null specificity (0/400) should not be conflated with perfect detection of technical factors: the separate recorded detection-sensitivity summary was 98/400 (24.5%) under its own definition.

**Fig. 2.**
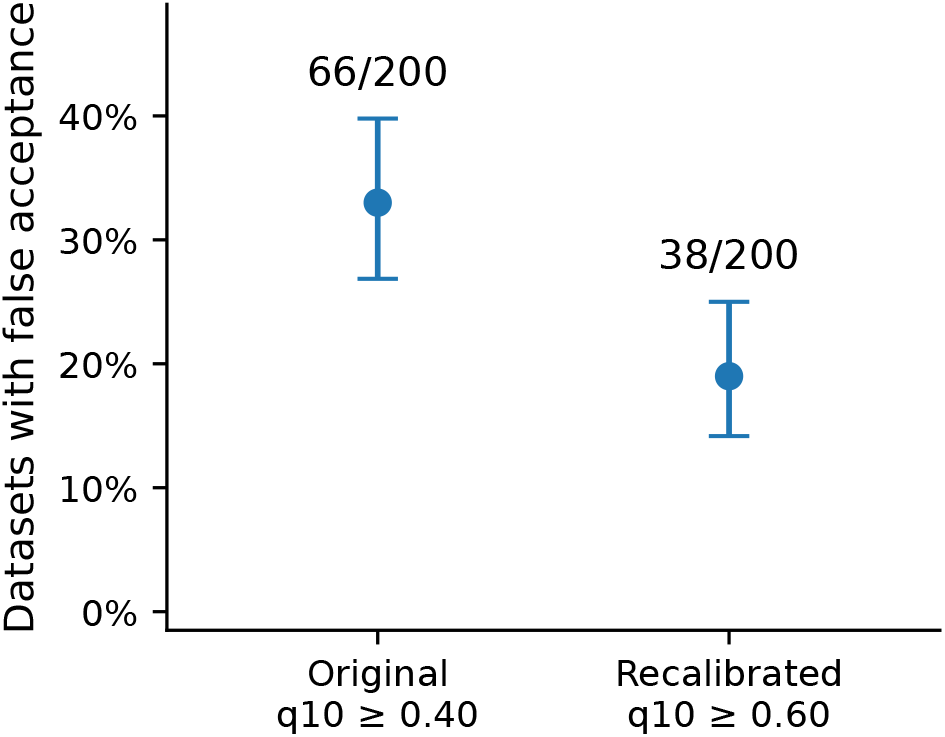
Fresh validation on 200 paired correlated-private datasets. Original and candidate rules were evaluated from the same fixed fit summaries. Marginal 95% Wilson bars are distinct from the paired reduction confidence interval.

### C. TARGET-OS: utility favored the simpler representation

Outcome-blind selection in 78 TARGET-OS patients favored V1 rather than the full V4 representation. In a read-only decomposition of the previously fixed outer-fold utility, V4 minus V1 was −0.020486 on average. The reconstruction component was −0.109115 (weighted contribution −0.032734), partly offset by adjusted cross-view retrieval +0.022468 (weighted +0.006740), missing-view score +0.007545 (weighted +0.001886), and modality balance +0.024147 (weighted +0.003622). This arithmetic accounts for the selected utility contrast; it does not estimate independent causal contributions of the training components. It also does not imply that *q*_0.10_ retention was applied to TARGET-selected V1.

In a post-freeze analysis of five full-data V1 coordinates, one of four preselected molecular programs, M34, exceeded matched random-program controls (maximum |*ρ*_*s*_| = 0.511 across coordinates, four-test BH-adjusted *q* = 0.004; 1,000 control panels). M40, M11, and M24 did not pass the same criterion (*q* = 0.172, 0.948, and 0.404). Technical covariate status was *unknown* and V1 coordinate-level resampling stability was not established. The association is therefore descriptive, not validation of a retained biological factor.

### D. TCGA external applications: task-dependent trade-offs

Table IV summarizes fixed-procedure results from five held-out folds. Relative to V1, the pre-retention V4 representation achieved better adjusted cross-view retrieval and lower missing-view error in both cohorts, but higher ordinary reconstruction error. The PCA baseline attained the lowest ordinary reconstruction NMSE in both. On SARC, PCA also had lower omitted-view NMSE than V4; in KIRC, V4’s advantage over PCA for omitted-view NMSE was small. These comparisons change multiple architecture components together and do not isolate a mechanism. Across both cohorts, all 20 neural epoch-selection runs per cohort reached the 400-epoch budget; numerical convergence is not established.

**TABLE III.** Audited OUTCOME-BLINDTARGET-OS COMPONENT DECOMPOSITION. Entries ARE V4 MINUS V1; POSITIVE COMPONENT CHANGES FAVOR V4, BUT THE FIXED TOTAL UTILITY FAVORS V1. These ARE POST-RESULT DESCRIPTIONS, NOT SEPARATE CAUSAL EFFECTS.

| TARGET utility component | Score diff. | Weighted |
| --- | --- | --- |
| Reconstruction (0.30) | −0.109115 | −0.032734 |
| Adjusted retrieval (0.30) | +0.022468 | +0.006740 |
| Missing view (0.25) | +0.007545 | +0.001886 |
| Modality balance (0.15) | +0.024147 | +0.003622 |
| **Aggregate utility** | — | **−0.020486** |

**TABLE IV.** External HELD-OUT FOLD MEANS (5 FOLDS; NEURAL VALUES AVERAGED OVER TWO SEEDS WITHIN FOLD). LOWER NMSE IS BETTER; LARGER ADJUSTED MRR IS BETTER. V4 IS EVALUATED *before* FACTOR RETENTION.

| Cohort | Model | Rec. NMSE | Missing NMSE | Adj. MRR |
| --- | --- | --- | --- | --- |
| SARC | PCA ($K = 10$) | **0.5845** | **0.7867** | 0.1820 |
|  | V1 | 0.7216 | 0.8643 | 0.1512 |
|  | V4 (pre-ret.) | 0.7322 | 0.8192 | **0.2113** |
| KIRC | PCA ($K = 10$) | **0.4391** | 0.8941 | 0.0706 |
|  | V1 | 0.7129 | 0.9834 | 0.0501 |
|  | V4 (pre-ret.) | 0.7270 | **0.8888** | **0.1198** |

## IV. Discussion AND Limitations

Three conclusions follow from these distinct experiments. First, retained latent factors cannot be equated with generator-designated biological factors solely because they recur across resampling fits. A structured null can contain real cross-view dependence and still fail the study’s chosen biological-shared truth criterion; the classifier and threshold results are conditional on this truth definition. This is an *identifiability and interpretation boundary*, not proof that the algorithm discovered an intrinsically meaningless direction.

Second, the prospective lower-tail threshold provides a measured improvement but not a universal solution. Several exploratory thresholds could meet recovery criteria, the selected 0.60 value was a trade-off rather than an optimum, 19% residual error remained, and exact modality-support attribution failed its historical test. The decoder-weight notion of support is parameterization-dependent. Neither the null results nor the single-family prospective simulation justify biological false-discovery guarantees across settings or independent datasets.

Third, the human-cancer applications support claims about *representations*, not factor retention. TARGET preferred V1 under an outcome-blind composite; SARC and KIRC show task-dependent gains and losses for pre-retention V4, not uniformly better performance. These cohorts differ in cancer type, sample size, and preprocessing, and their reconstruction metrics are not interchangeable with TARGET utility. No clinical outcome-prediction benefit is claimed. The observed TARGET program alignment is confined to one program under limited technical metadata, and two-seed agreement in external fits cannot replace full stability refitting.

Limitations include simulation-family dependence, a known failure of modality support, restricted *n* = 62 correlated-private stress tests, non-causal exploratory factor diagnostics, unavailable ground-truth factors in real data, a budget-limited external neural training protocol, and non-equivalent comparator capacities. We did not demonstrate that the recalibrated retention rule transfers to MOFA, JIVE/AJIVE, TARGET-selected V1, or external TCGA fits. These boundaries favor separate assessment of representation utility, retention, subspace recovery, and biological labeling rather than a single performance ranking.

## V. Conclusion

SABLE-MO supplies a controlled setting to test the gap between repeatable latent structure and justified factor retention. In a structured null, the original procedure showed failure that an independent-null control did not reveal. A prospectively fixed stricter rule reduced but did not eliminate that error. Across three cancer cohorts, representation trade-offs varied by task and did not establish biological validity of retained factors. The supported conclusion is a narrower methodological one: claims about latent factors require distinct evidence for reproducibility, operating characteristics, modality attribution, and biological interpretation.

## Code and Data availability

Source code, simulation contracts, result locks, and selected output summaries are in the public SABLE-MO repository (snapshot commit `2f8e13e1d63a`). Public TARGET/TCGA input manifests and acquisition procedures are documented there; a clean Git checkout is not claimed to contain every molecular matrix, checkpoint, or generated shard.

## Acknowledgment AND AI Assistance Disclosure

ChatGPT (OpenAI) assisted with drafting text in Sections I–V, and preparation of displayed figures from archived study summaries. Scientific values in this manuscript are drawn from the archived study records; authors are responsible for checking every claim, source and final wording before submission. No neural models were trained for the preparation of this manuscript.

## References

[1] E. F. Lock, K. A. Hoadley, J. S. Marron, and A. B. Nobel, “Joint and individual variation explained (JIVE) for integrated analysis of multiple data types,” The Annals of Applied Statistics, vol. 7, no. 1, pp. 523–542, 2013. [Online]. Available: 10.1214/12-AOAS597

[2] Q. Feng, M. Jiang, J. Hannig, and J. S. Marron, “Angle-based joint and individual variation explained,” Journal of Multivariate Analysis, vol. 166, pp. 241–265, 2018. [Online]. Available: 10.1016/j.jmva.2018.03.008

[3] R. Argelaguet, B. Velten, D. Arnol, S. Dietrich, T. Zenz, J. C. Marioni, F. Buettner, W. Huber, and O. Stegle, “Multi-omics factor analysis—a framework for unsupervised integration of multi-omics data sets,” Molecular Systems Biology, vol. 14, no. 6, p. e8124, 2018. [Online]. Available: 10.15252/msb.20178124

[4] S. D. McCabe, D.-Y. Lin, and M. I. Love, “Consistency and overfitting of multi-omics methods on experimental data,” Briefings in Bioinformatics, vol. 21, no. 4, pp. 1277–1284, 2020. [Online]. Available: 10.1093/bib/bbz070

[5] N. Meinshausen and P. Bühlmann, “Stability selection,” Journal of the Royal Statistical Society: Series B (Statistical Methodology), vol. 72, no. 4, pp. 417–473, 2010. [Online]. Available: 10.1111/j.1467-9868.2010.00740.x

[6] J. L. Ba, J. R. Kiros, and G. E. Hinton, “Layer normalization,” 1607.06450, 2016. [Online]. Available: https://arxiv.org/abs/1607.06450

[7] A. van den Oord, Y. Li, and O. Vinyals, “Representation learning with contrastive predictive coding,” 1807.03748, 2018. [Online]. Available: https://arxiv.org/abs/1807.03748

[8] I. Loshchilov and F. Hutter, “Decoupled weight decay regularization,” in International Conference on Learning Representations, 2019. [Online]. Available: https://openreview.net/forum?id=Bkg6RiCqY7

[9] P. H. Schönemann, “A generalized solution of the orthogonal Procrustes problem,” Psychometrika, vol. 31, no. 1, pp. 1–10, 1966. [Online]. Available: 10.1007/BF02289451

[10] H. W. Kuhn, “The Hungarian method for the assignment problem,” Naval Research Logistics Quarterly, vol. 2, no. 1–2, pp. 83–97, 1955. [Online]. Available: 10.1002/nav.3800020109

[11] E. B. Wilson, “Probable inference, the law of succession, and statistical inference,” Journal of the American Statistical Association, vol. 22, no. 158, pp. 209–212, 1927. [Online]. Available: 10.1080/01621459.1927.10502953

[12] Q. McNemar, “Note on the sampling error of the difference between correlated proportions or percentages,” Psychometrika, vol. 12, no. 2, pp. 153–157, 1947. [Online]. Available: 10.1007/BF02295996

[13] T. P. Morris, I. R. White, and M. J. Crowther, “Using simulation studies to evaluate statistical methods,” Statistics in Medicine, vol. 38, no. 11, pp. 2074–2102, 2019. [Online]. Available: 10.1002/sim.8086

[14] National Cancer Institute, “TARGET Osteosarcoma (TARGET-OS),” Genomic Data Commons, 2026, accessed October 2026. [Online]. Available: https://gdc.cancer.gov/content/target-os-publications-summary

[15] Cancer Genome Atlas Research Network, “Comprehensive and integrated genomic characterization of adult soft tissue sarcomas,” Cell, vol. 171, no. 4, pp. 950–965.e28, 2017.

[16] Cancer Genome Atlas Research Network, “Comprehensive molecular characterization of clear cell renal cell carcinoma,” Nature, vol. 499, pp. 43–49, 2013.

