## Supplemental for "Latent-Factor Stability Is Not Sufficient: Evaluating SABLE-MO in Simulated and Cancer Multi-Omics Data"

Elena Krikun and Abedalrhman Alkhateeb
Department of Computer Science, Lakehead University, Thunder Bay, Canada

---

**Scope.** This document expands the methods, audit logic, negative-control results, outcome-blind TARGET-OS model-selection analysis, full-data V1 sensitivity analyses, biological-program alignment, and external TCGA representation comparisons reported in the conference-length manuscript. The submitted BIBE 2026 main manuscript is intentionally left unchanged. This supplement does not introduce a new primary model-selection rule, reopen a frozen simulation gate, or use clinical outcomes to alter representation learning.

**Interpretive boundary.** Several analyses below are explicitly descriptive or post-result audits. Repeated cross-validation folds are not treated as independent experimental replicates. The stricter $q_{0.10} = 0.60$ retention rule was prospectively checked on fresh simulation seeds, but it is not transferred to TARGET-selected V1, to MOFA/JIVE/AJIVE, or to the external TCGA representations. Likewise, biological-program alignment is not treated as proof that an individual latent coordinate is a stable biological factor.

**Relationship to prior work by the authors.** Two earlier studies are directly relevant to the motivation for the present representation-learning analysis but are not used as evidence for its model-selection or retention decisions. The first introduced a dependency-aware Markov-Blanket framework for compact multimodal feature selection and cross-cohort reapplication in breast cancer [5]. The second separated assayability, literal model transport, within-cohort redevelopment, selection stability, and incremental prognostic performance in a TCGA-BRCA–METABRIC evaluation [6]. The present work addresses a different question: outcome-blind representation learning and reproducibility in small-sample cancer multi-omics, with TARGET-OS as the primary real-data application. Direct links: published Markov-Blanket study and cross-cohort multi-omics preprint.

**3 Supplementary Data and Code Notes** **12**

**4 Supplementary References** **12**

### 1 Supplementary Methods

#### 1.1 S1. Analysis stages, information barriers, and frozen decision logic

The project separated development, confirmatory simulation, real-data representation selection, full-data refitting, technical assessment, and biological interpretation. This separation was used to prevent favorable downstream results from feeding back into upstream representation choices. The major stages are summarized in Table 1.

**Table 1:** Major analysis stages and information-access boundaries.

| Stage | Role | Permitted information | Prohibited consequence |
|---|---|---|---|
| 05a–05e | Simulation development and method freeze | Simulated molecular data with known latent truth; prespecified model families, metrics, and retention rules | No TARGET outcome access; no post-result change to the original final gate |
| 05f | Historical final gate | Fresh/frozen simulation scenarios designated before gate evaluation | No threshold retuning after the gate result |
| 05y | Prospective retention-rule follow-up | Fresh, non-overlapping simulation seeds; same fitted summaries evaluated under original and candidate $q_{0.10}$ rules | Cannot rewrite the historical gate; cannot imply universal biological false-discovery control |
| 06b | TARGET representation benchmark | TARGET molecular inputs only; nested outcome-blind CV | No survival, outcome, clinical, or biology access; no downstream biology used for model selection |
| 06c | Full-data V1 refit | All 78 TARGET molecular profiles; frozen architecture/epochs/seeds | No post-fit seed selection; no outcome-guided refitting; no V1 use of the SABLE-MO V4 retention gate |
| 06d | Technical and transfer-boundary audits | Frozen V1 artifacts and available technical metadata | Missing metadata cannot be interpreted as evidence of no technical effect |
| 06e | Molecular-program alignment | Frozen V1 representation and preselected M34/M40/M11/M24 program scores | Biology cannot change model, seed, epoch, or representation family |
| 06e3–06e5a | Seed/subspace sensitivity and read-only audit | Three prospectively frozen V1 full-data fits and already-frozen program scores | No new seed-specific $p/q$ values; no model selection; no new stability threshold |

The governing distinction throughout the manuscript is between (i) representation utility, (ii) reproducibility/stability, (iii) factor-retention operating characteristics, (iv) modality-support attribution, and (v) biological interpretation. These quantities are related, but they are not interchangeable endpoints.

#### 1.2 S2. Representation ladder and the V3/V4 boundary

The neural development ladder increased complexity incrementally. The final TARGET-selected representation was V1, not V4. Table 2 records the interpretation used in the frozen audits.

A later read-only audit showed that V3 and V4 were numerically identical in the cross-family TARGET pre-retention benchmark: 75 corresponding rows were identical across utility, held-out reconstruction, adjusted cross-view MRR, missing-view reconstruction, modality-balance score, and selected epoch. Therefore, the TARGET benchmark does not support interpreting V4 as an independently worse representation architecture than V3. The meaningful comparison is V1 versus the V3/V4 pre-retention backbone; V4's additional scientific contribution is the separately characterized resampling alignment and retention procedure.

**Table 2:** Representation ladder used in development and TARGET benchmarking.

| Model | Complexity rank | Added representation component | Interpretation boundary |
|---|---|---|---|
| V0 | 6 | Block-balanced autoencoder baseline | Simpler neural integration baseline |
| V1 | 7 | Shared/private decomposition | Final TARGET-selected representation |
| V2 | 8 | Contrastive alignment added to V1 | Extra alignment complexity |
| V3 | 9 | Modality dropout and missing-view reconstruction added to V2 | More elaborate pre-retention representation backbone |
| V4 | 10 | V3 plus resampling-based factor alignment and stability/retention procedure | Distinctive V4 contribution is post-fit retention/stability governance, not a different pre-retention fit in the 06b cross-family benchmark |

#### 1.3 S3. Historical retention rule and factor correspondence

For each dataset, a reference fit was compared with 100 complete 80% subsample refits. Shared latent spaces were aligned using orthogonal Procrustes alignment [1]; factor identities were paired using Hungarian assignment [2]. If $r_{kb}$ denotes the absolute aligned correlation of reference factor $k$ with its matched factor in resample $b$, the inclusion frequency was

$$\pi_k = \frac{1}{100} \sum_{b=1}^{100} \mathbf{1}\{r_{kb} \geq 0.50\}. \tag{1}$$

The original and prospective candidate rules are reproduced in Table 3. Only the lower-decile threshold changed in the prospective follow-up.

**Table 3:** Factor-retention conditions. All Boolean criteria must be satisfied.

| Criterion | Original | Candidate | Role |
|---|---|---|---|
| Median aligned correlation | $\geq 0.70$ | $\geq 0.70$ | Central resampling reproducibility |
| Lower-decile aligned correlation, $q_{0.10}$ | $\geq 0.40$ | $\geq 0.60$ | Lower-tail reproducibility |
| Inclusion frequency | $\geq 0.80$ | $\geq 0.80$ | Fraction of resamples with correlation at least 0.50 |
| Factor-score variance | $\geq 10^{-4}$ | $\geq 10^{-4}$ | Excludes numerically collapsed factors |
| Supported modalities | $\geq 2$ | $\geq 2$ | Requires multi-view support |
| Per-view relative decoder energy defining support | $\geq 0.15$ | $\geq 0.15$ | Parameterization-specific support definition |

The support-energy quantity is tied to the frozen decoder parameterization. It is not interpreted as a universal feature-importance measure. Rotational subspace recovery is reported separately because stable subspaces can exist even when individual axes rotate.

##### 1.4 S4. Known-truth simulation families and denominators

The simulator generated three-view RNA-like, methylation-like, and copy-number-like data with biological shared factors, modality-private factors, technical factors, feature offsets, and noise. The generic form was

$$X^{(v)} = SA_v^\top + P^{(v)}B_v^\top + TC_v^\top + E^{(v)}. \tag{2}$$

The historical confirmatory registry spanned 24 scenario families and 3,100 final-gate datasets. The specific null failure discussed in the manuscript occurred in a correlated-private mechanism at $n = 62$, in which no factor was designated as biological shared truth but modality-private scores remained cross-view dependent. For that family,

$$P^{(v)} = \operatorname{stdcol}\left(\sqrt{0.60}\,U + \sqrt{0.40}\,E_v\right). \tag{3}$$

The following denominators were intentionally kept separate:

- **Dataset-level false acceptance:** a dataset counts once if at least one retained nontechnical factor fails to match any registered biological-shared truth.
- **Truth-factor recovery:** denominator is the number of true biological shared-factor opportunities, not the number of datasets.
- **Exact modality-support attribution:** denominator is restricted to truth-matched retained biological factors for which exact generating support is assessable.
- **Technical false biological acceptance:** denominator is the prespecified set of technical-factor opportunities, distinct from technical-factor detection sensitivity.

Wilson intervals were used for marginal proportions [3]. In the fresh prospective paired validation, the original and candidate rules were evaluated on the same fitted summaries; discordance was therefore evaluated with an exact McNemar comparison [4], and the absolute paired reduction used a dataset bootstrap.

##### 1.5 S5. Prospective $q_{0.10}$ follow-up design

The historical correlated-private failure prompted an exploratory diagnostic stage. That stage was not itself confirmatory. Candidate lower-decile thresholds were examined only on historical data. The value $q_{0.10} = 0.60$ was then fixed before creating a new seed bank containing 200 correlated-private datasets and 200 standard-signal datasets with four true shared factors each. Random seeds did not overlap the historical bank. Both the original and candidate rules were evaluated from the same frozen per-dataset fit summaries, so no duplicate model fitting was required for the rule comparison.

The fresh validation had two prespecified qualitative requirements: the candidate should reduce the correlated-private dataset-level false-acceptance rate and should preserve adequate true shared-factor recovery. The analysis was a mechanism-specific validation within the same simulator family, not external biological validation.

##### 1.6 S6. TARGET-OS outcome-blind nested representation benchmark

TARGET-OS contained 78 patients with matched RNA expression, DNA methylation, and gene-level copy number. Representation selection used five repeats of five-fold outer cross-validation. Each outer training set contained an inner four-fold model-selection procedure. All preprocessing and model fitting remained within the applicable training partition.

The outcome-blind TARGET utility was

$$U_{\text{TARGET}} = 0.30H_{\text{rec}} + 0.30M_{\text{retrieval}} + 0.25H_{\text{missing}} + 0.15B_{\text{modality}}, \tag{4}$$

where the four terms summarize held-out reconstruction, chance-adjusted same-patient cross-view retrieval, missing-view reconstruction, and modality balance. No survival outcome, clinical variable, biological-program score, or downstream interpretation entered this utility.

After prespecified failure gates, the selection rule identified the best inner-CV mean, computed the one-standard-error cutoff using the standard error of that best model, formed the eligible set above the cutoff, and selected the lowest-complexity eligible method. The 25 outer splits are repeated cross-validation splits, not independent biological replicates; all counts and differences below are therefore descriptive unless explicitly stated otherwise.

#### 1.7 S7. Full-data V1 refit and seed policy

Following selection of V1, a separate full-data refit contract froze the compact/weak profile, RNA/methylation/CNV feature budgets of 5,000/10,000/5,000, up to ten PCA dimensions per modality, five shared dimensions, two private dimensions per modality, and 400 fixed epochs. The primary full-data seed was 1103. Seeds 2207 and 3301 were prospectively designated as reproducibility-only fits and could not replace the primary representation based on downstream results.

The final full-data representation had shape $78 \times 5$ for each seed. The V4/SABLE-MO $q_{0.10}$ retention gate was explicitly not transferred to V1. Likewise, no post-fit ensemble or seed averaging was permitted unless separately frozen.

#### 1.8 S8. V1 seed/subspace sensitivity

Because raw latent coordinates can differ by sign, permutation, or rotation even when their subspaces agree, the seed-sensitivity audit compared the three frozen V1 full-data fits at the subspace level. For each pair, centered latent matrices were converted to rank-preserving orthonormal bases. The singular values of $Q_A^\top Q_B$ were reported as canonical correlations; principal angles were their inverse-cosine transforms. The pairwise mean canonical correlation was summarized descriptively. Orthogonal Procrustes alignment followed by Hungarian matching was reported as a complementary coordinate-level diagnostic.

No V1 stability threshold had been prospectively frozen, so these quantities do not generate a PASS/FAIL decision. They are sensitivity diagnostics only.

#### 1.9 S9. Frozen biological-program alignment

Four preselected molecular programs were carried forward from the earlier molecular-transport analysis: M34 (162 genes), M40 (111), M11 (7), and M24 (7). Within the exact 78 TARGET cases, each frozen signed program score was standardized with sample standard deviation (ddof = 1). Program-gene coverage was 100% for the required frozen members.

For primary seed 1103, the biological-alignment statistic for each program was the maximum absolute Spearman correlation across the five raw frozen V1 shared coordinates. All 20 coordinate-program correlations were retained, and the maximizing coordinate was recorded. Program-specificity was evaluated against 1,000 matched random gene programs using a prespecified matched-control procedure. Multiplicity correction used Benjamini–Hochberg across the four programs.

This statistic is sign/permutation invariant at the maximizing-coordinate level, but it is not rotation invariant. Therefore, the later seed/subspace sensitivity did not recompute matched-random $p$-values for seeds 2207 or 3301. The correct interpretation is that M34 was matched-null-supported in the prospectively fixed primary seed, with similar maximum-alignment magnitudes observed in the two prospectively designated reproducibility fits.

#### 1.10 S10. External TCGA representation comparisons

TCGA-SARC ($n = 250$) and TCGA-KIRC ($n = 304$) were fitted independently within cohort. The comparison involved budgeted block PCA ($K = 10$), V1, and the pre-retention V4 representation. Five outer folds were used; neural fold summaries averaged two fixed seeds. Ordinary reconstruction and omitted-view errors are normalized mean squared errors (lower is better). Chance-adjusted cross-view MRR is higher-is-better.

These experiments do not validate the V4 retention rule. They compare representation behavior before retention and therefore support only task-specific representation statements.

### 2 Supplementary Results

#### 2.1 S11. Historical final-gate endpoint audit

Table 4 expands the abbreviated gate summary in the main manuscript. D01–D07 are intentionally heterogeneous: they do not share a common denominator or interpretation.

**Table 4:** Historical final-gate outcomes under the original retention rule.

| ID | Endpoint | Result | Interpretation |
|---|---|---|---|
| D01 | Pooled biological-null dataset-level false acceptance | 70/1000 = 7.0%; Wilson 95% CI approximately 5.58–8.75% | Passed the prespecified pooled criterion, but all failures came from one structured-null family |
| D02 | Standard-signal true shared-factor recovery | 371/400 = 92.75%; Wilson 95% CI approximately 89.78–94.90% | Passed recovery criterion |
| D03 | Standard-signal false-negative rate | 29/400 = 7.25%; Wilson 95% CI approximately 5.10–10.22% | Complementary recovery endpoint |
| D04 | Rotational subspace recovery | mean canonical-correlation summary 0.828986 | Passed the 0.80 subspace criterion; does not imply stable individual axes |
| D05 | Exact generating-modality support among eligible retained biological factors | 81/120 = 67.5% | Failed the prespecified 80% criterion |
| D06 | False biological acceptance per true technical-factor opportunity | 0/400 | Passed specificity criterion; separate technical-factor detection sensitivity was only 98/400 |
| D07 | Process integrity / no prohibited retuning | PASS | Governance endpoint, not a predictive-performance statistic |

The key historical observation is that a passing pooled D01 result concealed complete concentration of the 70 false-acceptance events in the correlated-private null. At matched $n = 62$, the independent-null subset had 0/200 events whereas the correlated-private null had 70/200. The groups are separate generative mechanisms and are not paired observations.

#### 2.2 S12. Exploratory mechanism diagnostics and fresh prospective validation

Removing the support-count or factor-variance requirements did not change the historical matched-$n$ correlated-private false-acceptance count of 70/200. A grouped out-of-fold exploratory classifier using median, lower-decile, and inclusion summaries achieved AUC 0.967 (bootstrap 95% interval 0.950–0.981) for distinguishing retained false factors from retained standard-signal factors. Because scenario and truth status are coupled in this comparison, it is not interpreted as a general biological-factor classifier. A non-retained scenario-label anchor had AUC 0.792, and realized private-score correlation weakly discriminated false-acceptance events (AUC 0.579; interval included 0.5).

Historical threshold slicing produced multiple trade-offs, demonstrating that $q_{0.10} = 0.60$ was not mathematically unique. With the median and inclusion thresholds unchanged, the historical correlated-private false-acceptance counts and standard truth recoveries were 41/200 and 368/400 at $q_{0.10} = 0.60$, versus 18/200 and 355/400 at $q_{0.10} = 0.65$.

The residual 19% false-acceptance rate is substantial. The follow-up therefore supports a narrower statement: increasing lower-tail reproducibility reduced error in this prespecified structured-null mechanism without materially changing recovery in the paired fresh bank.

**Table 5:** Fresh paired validation of the prospectively fixed $q_{0.10} = 0.60$ candidate.

| Endpoint | Original | Candidate | Paired interpretation |
|---|---|---|---|
| Correlated-private dataset-level false acceptance | $66/200 = 33\%$ | $38/200 = 19\%$ | Absolute paired reduction 14 percentage points; dataset-bootstrap 95% CI 9.5–19.0 |
| Discordant correlated-private datasets | 28 candidate improvements | 0 reversals | Exact two-sided McNemar $p = 7.45 \times 10^{-9}$ |
| Standard-signal true-factor recovery | $755/800 = 94.375\%$ | $752/800 = 94.0\%$ | Small absolute reduction (0.375 percentage points) |
| Historical D05 modality-support result | 81/120 | unchanged | Candidate $q_{0.10}$ follow-up did not reopen the historical support-attribution failure |

#### 2.3 S13. TARGET one-standard-error selection audit

Across the 25 outer splits, V1 was selected in 17 and V0 in 8. V1 had the highest inner-CV mean in 23/25 splits; V2 and V3 each had the best inner mean once, and V4 never uniquely appeared as the recorded best model. The one-standard-error acceptance counts were V0: 8/25, V1: 25/25, V2: 17/25, V3: 16/25, and V4: 16/25. Block PCA and all MOFA variants were outside the one-standard-error acceptance set in all 25 splits.

Because V0 is simpler than V1, the exact implication of the selection logic is especially useful: in every one of the 17 splits where V1 was ultimately selected, V0 was below the one-standard-error cutoff. This is stronger and more precise than describing the difference as a generic "greater than one SE" effect, which is not the rule being applied.

**Table 6:** TARGET nested-CV selection audit across 25 repeated outer splits.

| Model | Selected by one-SE rule | Best inner mean |
|---|---|---|
| BLOCK_PCA | 0 | 0 |
| MOFA $K = 5/10/15/20$ | 0 each | 0 each |
| V0 | 8 | 0 |
| V1 | 17 | 23 |
| V2 | 0 | 1 |
| V3 | 0 | 1 |
| V4 | 0 | 0 |

#### 2.4 S14. Descriptive held-out utility margins for TARGET V1

The frozen outer-CV results were audited after averaging seeds within each outer split. Table 7 summarizes descriptive V1-minus-comparator utility differences. These 25 folds arise from repeated CV and are not treated as 25 independent experiments; no new inferential $p$-values or confidence intervals were introduced.

**Table 7:** Descriptive V1-minus-comparator held-out utility differences across 25 outer splits.

| Comparator | Mean $\Delta U$ | SD | Median | V1 higher | SD of 5 repeat means |
|---|---|---|---|---|---|
| BLOCK_PCA | 0.114390 | 0.048375 | 0.117078 | 25/25 | 0.006816 |
| MOFA $K = 5$ | 0.131270 | 0.037684 | 0.132725 | 25/25 | 0.015987 |
| MOFA $K = 10$ | 0.074476 | 0.042181 | 0.078690 | 24/25 | 0.011885 |
| MOFA $K = 15$ | 0.056738 | 0.047790 | 0.064594 | 23/25 | 0.009505 |
| MOFA $K = 20$ | 0.051009 | 0.045073 | 0.057370 | 21/25 | 0.014012 |
| V0 | 0.028295 | 0.013335 | 0.028692 | 24/25 | 0.006465 |
| V2 | 0.018290 | 0.022222 | 0.016357 | 19/25 | 0.011242 |
| V3 | 0.020486 | 0.022354 | 0.018900 | 21/25 | 0.007795 |
| V4 | 0.020486 | 0.022354 | 0.018900 | 21/25 | 0.007795 |

V1 also showed lower seed sensitivity than the V3/V4 pre-retention backbone in the frozen outer-CV audit: the mean within-split seed SD of utility was 0.013744 for V1 versus 0.028444 for V4, and V4 had the larger seed SD in 88% of splits. Between-split SD of the seed-mean utility was 0.032159 for V1 and 0.038009 for V4. Because V3 and V4 were identical in this benchmark, these quantities characterize the V3/V4 pre-retention backbone rather than a V4-specific retention effect.

#### 2.5 S15. Exact V3/V4 pre-retention identity check

The cross-family TARGET result table contained 75 V3 rows and 75 V4 rows with identical split/seed keys. Exact row-by-row comparison found zero differences in utility, held-out reconstruction, adjusted cross-view MRR, missing-view reconstruction, modality-balance score, or selected epoch. Maximum absolute difference was zero for each metric. This follows the frozen method registry: V4 adds resampling-based factor alignment and the stability/retention gate to V3, while the cross-family benchmark did not apply the V4 post-fit retention gate.

Consequently, "V4 selected 0/25" is not negative evidence against the retention/stability procedure itself. V4 had the higher complexity rank, and an identical pre-retention utility to V3 is structurally disfavored by the one-standard-error simplicity rule.

#### 2.6 S16. Full-data V1 seed/subspace sensitivity

The final primary V1 fit remained seed 1103. Two additional full-data fits at seeds 2207 and 3301 were prospectively designated for reproducibility sensitivity only. Pairwise five-dimensional subspace agreement was moderate rather than high (Table 8).

**Table 8:** Pairwise agreement among three frozen full-data V1 fits.

| Seed pair | Mean canonical corr. | Minimum canonical corr. | Maximum principal angle | Normalized Procrustes residual |
|---|---|---|---|---|
| 1103–2207 | 0.556543 | 0.265451 | 74.606° | 0.849762 |
| 1103–3301 | 0.496533 | 0.080458 | 85.385° | 0.944832 |
| 2207–3301 | 0.618488 | 0.318616 | 71.421° | 0.839422 |

The median of the three pairwise mean canonical correlations was 0.556543. This statistic was retained only as a descriptive compatibility quantity with the earlier generic seed-aligned subspace metric. No V1 threshold or stability gate was defined for it. In particular, the 1103–3301 comparison included a canonical direction with correlation approximately 0.08 (principal angle approximately 85.4°), indicating that the independent full-data fits are not related merely by sign, permutation, or a near-perfect orthogonal rotation.

#### 2.7 S17. Primary biological-program alignment in V1 seed 1103

Table 9 gives the frozen primary biological-alignment results. The maximizing statistic was maximum absolute Spearman correlation across the five V1 shared coordinates. The matched-random empirical probability used 1,000 matched gene programs; BH correction was applied over the four prespecified programs.

**Table 9:** Primary seed-1103 molecular-program alignment.

| Program | Max $\lvert\rho_s\rvert$ | Best coordinate | Signed $\rho_s$ | BH $q$ | Interpretation |
|---|---|---|---|---|---|
| M34 | 0.510667 | Shared 04 | −0.510667 | 0.003996 | Matched-null-supported in the prospectively fixed primary fit |
| M40 | 0.394605 | Shared 01 | +0.394605 | 0.171828 | Not supported after four-program correction |
| M11 | 0.150976 | Shared 04 | −0.150976 | 0.948052 | Not supported |
| M24 | 0.329304 | Shared 05 | −0.329304 | 0.403596 | Not supported |

Technical covariate status for these TARGET factors was *unknown*: the technical-metadata preflight did not identify a strict usable covariate set. Unknown status cannot be converted into a "nontechnical" label. In addition, coordinate-level stability had not been established when the primary biology test was frozen. Accordingly, the M34 result is described as alignment of a frozen V1 shared coordinate with a prespecified molecular program, specific relative to matched random programs, rather than discovery of a stable biological factor.

#### 2.8 S18. Program-level sensitivity across the frozen reproducibility fits

The two reproducibility-only V1 fits were evaluated without transferring the primary matched-random $p/q$ values. Table 10 shows the raw maximum absolute Spearman magnitudes across seeds.

**Table 10:** Maximum absolute program-coordinate Spearman correlation across three prospectively frozen V1 full-data fits. Only seed 1103 has the primary matched-random $p/q$ analysis.

| Program | Seed 1103 | Seed 2207 | Seed 3301 | Range |
|---|---|---|---|---|
| M34 | 0.510667 | 0.453104 | 0.448830 | 0.061837 |
| M40 | 0.394605 | 0.294478 | 0.490939 | 0.196462 |
| M11 | 0.150976 | 0.203657 | 0.134056 | 0.069601 |
| M24 | 0.329304 | 0.192251 | 0.246273 | 0.137053 |

M34's alignment magnitude recurred across all three fits despite the only moderate subspace agreement, which argues against an interpretation in which the M34 alignment exists only in the primary seed. However, the two sensitivity fits do not receive new significance labels. M40 illustrates why seed sensitivity cannot be converted into post-hoc program selection: at seed 3301 its raw maximum alignment exceeds M34's, yet the frozen primary testing family remains the seed-1103 analysis.

Exploratory Procrustes-oriented sensitivities for the two reproducibility fits were: M34, 0.488436 (2207) and 0.381429 (3301); M40, 0.374372 and 0.314230; M11, 0.219388 and 0.132007; M24, 0.209499 and

0.260259. Procrustes orientation did not fully restore the primary M34 magnitude, particularly for seed 3301, which is consistent with genuine seed-dependent changes in the estimated five-dimensional subspace.

##### 2.9 S19. External TCGA representation results

The external cohort means are reproduced in Table 11 for completeness. V4 improved adjusted retrieval and missing-view reconstruction relative to V1 in both cohorts, while worsening ordinary reconstruction. Block PCA produced the lowest ordinary reconstruction NMSE in both cohorts. These mixed results motivate task-specific rather than universal superiority claims.

**Table 11:** External pre-retention representation comparison. Five held-out folds; neural entries average two fixed seeds within fold.

| Cohort | Model | Reconstruction NMSE | Missing-view NMSE | Adjusted MRR |
|---|---|---|---|---|
| SARC | Block PCA ($K = 10$) | 0.5845 | 0.7867 | 0.1820 |
| SARC | V1 | 0.7216 | 0.8643 | 0.1512 |
| SARC | V4 (pre-retention) | 0.7322 | 0.8192 | 0.2113 |
| KIRC | Block PCA ($K = 10$) | 0.4391 | 0.8941 | 0.0706 |
| KIRC | V1 | 0.7129 | 0.9834 | 0.0501 |
| KIRC | V4 (pre-retention) | 0.7270 | 0.8888 | 0.1198 |

All 20 neural epoch-selection runs per cohort reached the 400-epoch budget, so numerical convergence is not established. The external-cohort results therefore remain held-out representation comparisons under a fixed budget rather than evidence that one architecture universally converges better.

##### 2.10 S20. Reproducibility/provenance checkpoints

Table 12 lists selected immutable artifacts that anchor the analyses expanded above. The purpose of the table is provenance rather than reader-facing interpretation; scientific conclusions should be drawn from the reported estimands, not from the presence of hashes alone.

**Table 12:** Selected frozen artifacts and SHA-256 identifiers.

| Artifact | SHA-256 | Role |
|---|---|---|
| 06b authorized TARGET benchmark runner | `4f69680a2d26d6c8e4bd5b337a5e94e6cdc36d149ea6a4b9af547b3855a93eab` | Outcome-blind cross-family TARGET benchmark |
| 06b benchmark result lock | `2730c783f1a74f7aa58030d1ee03f324f1d50c4830c72e4229a15e41b60dc121` | Freezes TARGET benchmark outputs |
| 06c full-data V1 refit contract | `8ab8516e30964f220d0f1204ca892e29ca337691e06cbcbf29315e1cb6aec2e5` | Freezes primary full-data V1 configuration |
| 06c final authorized V1 runner | `006f718de8b0a1f768f8b4b7c35e4951719970f4d91ef43fa75541c74c8dc3bb` | Full-data primary/reproducibility fits |
| 06c3 full-data result lock | `b23d4c630cffca3bc5b1c44328128d4da46028b8db8cb070e6be93d9d0795be5` | Freezes three full-data V1 results |
| Primary seed-1103 V1 latent array | `c05bf08017ad8544e1e4ce0c8f58f086060dd7a6a0a68ed7f8d5a6723768a95f` | $78 \times 5$ primary shared latent matrix |
| 06e primary biological summary | `e4e68f5f21b2a54db4d0524fdaba265c42972c0fb6654036d9116f5ed3ca8370` | Primary M34/M40/M11/M24 matched-null analysis |
| 06e3 seed-sensitivity contract | `cbac52ec634e566ccee070ec0116577388a23d4ceefd881fbe761b55a39e40a6` | Freezes post-result sensitivity scope |
| 06e4 seed/subspace runner | `97a8af048e38c16000f27a078ea976af1c894a21850ac2924a05e3b8c16cee86` | Computes three-seed subspace/program sensitivity |
| 06e5a v2 read-only audit script | `72bb437d7794b62bb3c332e4bf21a2f95d56beb69cd8cc8722e280faa065ffc2` | Independent replay/audit of frozen seed-sensitivity outputs |

##### 2.11 S21. Interpretation rules carried into the preprint

The supplement uses the following wording constraints to keep the evidence hierarchy explicit:

- The historical pooled null gate *passed*, but a specific correlated-private mechanism failed substantially.
- The $q_{0.10} = 0.60$ candidate *reduced* this mechanism-specific error on fresh seeds; it did not eliminate error or establish distribution-free error control.
- V1 was selected in TARGET by an outcome-blind one-standard-error rule. The V4 retention threshold was not transferred to V1.
- V3 and V4 were identical as pre-retention representations in the cross-family TARGET benchmark; therefore the benchmark cannot attribute their seed sensitivity specifically to the V4 retention procedure.
- The V1 five-dimensional full-data subspace was materially seed-dependent. There is no prospectively defined V1 stability PASS/FAIL threshold.
- M34 was matched-null-supported in the fixed primary seed; comparable effect magnitudes in the two reproducibility fits are sensitivity evidence, not two additional significance tests.
- Technical status for TARGET V1 biological interpretation remains unknown, not clean or nontechnical.
- External SARC/KIRC results compare representation tasks before retention and do not validate factor retention.

### 3 Supplementary Data and Code Notes

The reproducibility archive contains versioned contracts, seed registries, result locks, scripts, and selected result summaries. Large molecular matrices, model checkpoints, and all generated simulation shards are not assumed to be present in a minimal repository checkout. Public cohort acquisition manifests and processing instructions are maintained separately from outcome variables. The numerical results reported in this supplement were drawn from frozen study artifacts and read-only audit summaries; no clinical outcome was used to select the TARGET representation or to alter the biological-alignment analysis.

### 4 Supplementary References

[1] P. H. Schoenemann, “A generalized solution of the orthogonal Procrustes problem,” *Psychometrika*, vol. 31, no. 1, pp. 1–10, 1966.

[2] H. W. Kuhn, “The Hungarian method for the assignment problem,” *Naval Research Logistics Quarterly*, vol. 2, no. 1–2, pp. 83–97, 1955.

[3] E. B. Wilson, “Probable inference, the law of succession, and statistical inference,” *Journal of the American Statistical Association*, vol. 22, no. 158, pp. 209–212, 1927.

[4] Q. McNemar, “Note on the sampling error of the difference between correlated proportions or percentages,” *Psychometrika*, vol. 12, no. 2, pp. 153–157, 1947.

[5] E. Krikun and A. Alkhateeb, “A Markov Blanket-based framework for dependency-aware feature selection in multimodal breast cancer data,” *Network Modeling Analysis in Health Informatics and Bioinformatics*, vol. 15, art. 158, 2026. doi: 10.1007/s13721-026-00832-1.

[6] E. Krikun and A. Alkhateeb, “External Evaluation of Multi-Omics Prognostic Models Across TCGA-BRCA and METABRIC: Transportability, Stability, and Incremental Performance,” *bioRxiv*, 2026. doi: 10.64898/2026.09.18.752709.
